# Longitudinal Identification of Zebrafish Individuals by Deep Learning

**DOI:** 10.64898/2025.12.14.694189

**Authors:** Danying Cao, Cheng Guo, Yingyin Cheng, Wanting Zhang, Mijuan Shi, Xiao-Qin Xia

## Abstract

The zebrafish (*Danio rerio*) is a critical vertebrate model organism in biomedical research. Accurate identification and longitudinal tracking of individual fish within cohorts enable linking genotype to complex phenotypes, essential for elucidating the molecular mechanisms underlying disease pathogenesis and trait formation. Nevertheless, the establishment of reliable long-term individual identification remains a significant challenge, primarily due to their diminutive size and minimal interindividual morphological variations. We developed ESC-IDNet, a dual-stage deep learning cascade architecture deployed on the FishIndivID platform (http://bioinfo.ihb.ac.cn/fishindivid), enabling high-precision identification. Trained and tested on ∼300,000 images from 450 zebrafish (31-122 dpf), our lateral body-based identification maintained >95% accuracy with updates only every 20 days, significantly outperforming dorsal head identification (requiring ∼13-day updates). Both views achieved 100% accuracy in 1-2 day tasks. ESC-IDNet surpassed alternatives in segmentation, alignment, feature extraction, and feature matching. This framework provides a robust, transferable paradigm for individual identification in fish species.

## 1. Introduction

The zebrafish (*Danio rerio*) is a pivotal vertebrate model organism in biomedical research [1], whose utility in linking genetic underpinnings to complex phenotypic and behavioral outcomes heavily relies on the accurate longitudinal tracking of individuals. The ability to uniquely identify individual fish over time unlocks a wide range of critical applications. For instance, in high-throughput drug screening, it enables the correlation of long-term treatment responses with individual metabolic or behavioral histories, moving beyond population averages to uncover heterogeneous drug effects. In behavioral neuroscience, individual recognition is prerequisite for studying social hierarchy formation, learning and memory trajectories, and personality differences within shoals. Furthermore, in developmental genetics and disease modeling, it allows for the precise monitoring of phenotypic onset and progression—such as tumor growth or neurological decline—in specific genotypes, thereby bridging the gap between genotype and phenotype with high temporal resolution. However, the establishment of a reliable, long-term individual identification system for zebrafish remains a significant challenge, primarily due to their small size and minimal interindividual morphological variation.

Conventional marking methods, such as passive integrated transponder (PIT) tags, are impractical for small-bodied zebrafish [2, 3]. Visible implant elastomer (VIE) tags [4] and physical markers like fin-clipping [5] are invasive, inducing physiological stress that can compromise experimental integrity. While manual identification is an alternative[6], it is labor-intensive and error-prone, especially in large-scale studies. Consequently, non-invasive, automated identification based on morphological appearance presents an attractive solution. Computer vision and deep learning technologies, celebrated for their cost-effectiveness and high accuracy in biometric recognition [7], have indeed seen extensive application in fish morphology analysis for tasks with substantial inter-class variations, such as species classification [8], disease detection [9, 10], size measurement [11], and fry grading [12]. Yet, for the task of individual identification, this approach is fraught with difficulties. Naturally occurring features are often extremely subtle, difficult to formulate manually, and subject to change due to growth and environmental influences, which can compromise long-term recognition reliability.

In broader fish recognition research, automated individual identification has been more successfully applied to patterned species with distinctive markings, such as Atlantic salmon, tiger barb, epaulette shark, and so on [13–18]. These studies typically report high short-term accuracy (≥94%), but long-term performance (over months) exhibits significant volatility (70%–93%) due to ontogenetic changes [13–16]. For instance, ventral spot-based identification in Atlantic salmon dropped from 100% accuracy at 2 months to 70% after 6 months [14]. The highest reported long-term accuracy of 93% at 10-month intervals was achieved by selectively sampling the most distinctive individuals from a larger cohort [13], an approach that lacks generalizability. In contrast, research on non-patterned species (e.g., perch, carp, delta smelt) is limited and reports considerably lower long-term accuracy (≤50%) [19–21]. Most existing studies, whether on patterned or non-patterned species, are further constrained by limited sample sizes (typically 10-100 individuals) and short validation periods, failing to provide a robust and generalizable long-term solution.

As a patterned species, zebrafish stands to benefit from visual identification, but research in this domain is scarce. Existing literature demonstrates limited utility: one approach utilizing stripe pigmentation features with k-nearest neighbors (KNN) classification achieved 99% accuracy in single-day trials but was validated on only five fish [22]. Another study employing a CNN-ViT hybrid model extended the recognition interval to 3 weeks but required daily image updates to maintain ≥95% accuracy, also using a minimal validation set [23]. These limitations underscore a critical gap in the field—the lack of a method that balances high accuracy, long-term stability, and broad generalizability for individual zebrafish.

In order to address these challenges, this study conducted longitudinal imaging of 450 zebrafish from larval to adult stages, manually curated approximately 300,000 image frames, and developed an integrated zebrafish identification framework—ESC-IDNet—by combining deep learning and computer vision technologies (Figure 1). This framework achieves individual-level identification by calculating the similarity between zebrafish across different time points. The ESC-IDNet is comprised of two core modules: ESCAlignNet and IDNet. ESCAlignNet, an improved version of YOLO [24], employs a dual-region segmentation strategy (lateral body and dorsal head regions) with keypoint-driven pose verification to achieve precise segmentation and standardized alignment of image regions. The IDNet module is predominantly comprised of an IResNet backbone network that has been integrated with a dynamic ArcFace loss function. This integration facilitates highly accurate and temporally stable identification from standardized images. ESC-IDNet has been demonstrated to enhance recognition accuracy, attaining 100% short-term identification accuracy. When combined with an adaptive updating strategy, it has been demonstrated to maintain over 95% accuracy in long-term recognition. The methodology developed in this study offers a novel solution for individual zebrafish identification and provides valuable insights for research on other fish species. It has the potential for extensive utilization in scientific research involving experimental animals.

**Figure 1.**
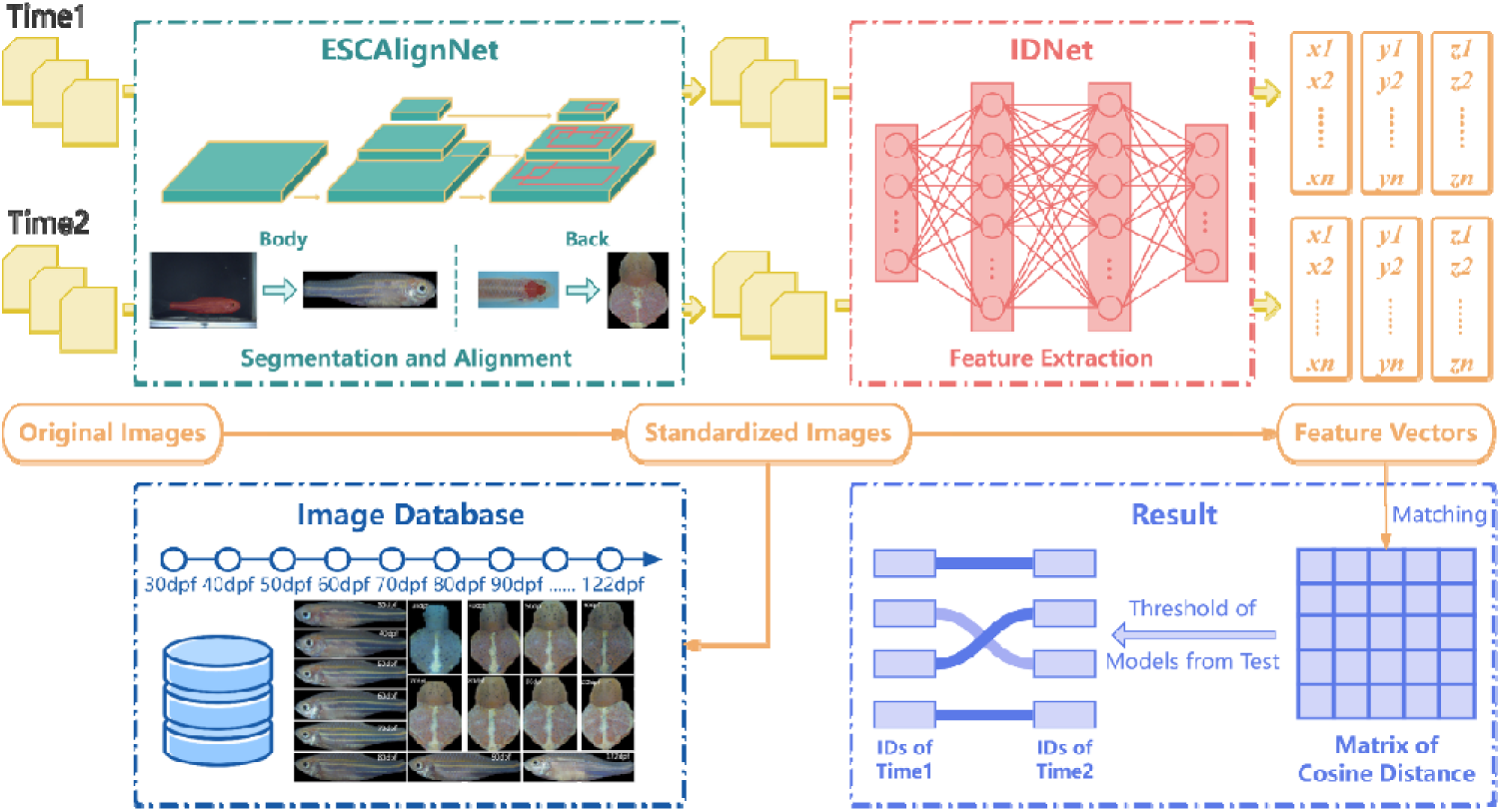
Overall process of zebrafish individual identification. ESCAlignNet performs target detection, segmentation and alignment standardization on images. IDNet performs feature extraction on images. The Image Database stores and registers standardized images for individual identification management. The result determines the individual identity based on the threshold and outputs it.

## 2. Materials and methods

### 2.1 Zebrafish breeding and image acquisition

#### 2.1.1 Zebrafish breeding

This study utilized zebrafish strains exhibiting diverse body coloration patterns, including wild-type (normal pigmentation), red, yellow, green, blue, purple, and transparent phenotypes (Supplementary Figure 1A). Two experimental groups (A and B) were randomly selected from the laboratory zebrafish population. The fish were fed Artemia nauplii twice daily, with the nauplii being freshly hatched. The fish were maintained under a controlled photoperiod of 10 hours of light and 14 hours of darkness. This photoperiod was selected to simulate natural diurnal cycles. Group A consisted of 150 zebrafish at 1-month post-fertilization (mpf), and no significant intergroup differences were observed in standard body length or weight. The group under consideration included 56 wild-type individuals and 94 fish exhibiting variant colorations. Each fish was housed individually in a transparent rearing chamber (10 × 10 × 10 cm), which was positioned within a recirculating aquatic system (RAS). A continuous flow of filtered and oxygenated water through apertures at both ends of each chamber ensured water quality maintenance. The water temperature was maintained at 27°C, and the pH level was kept at 7.4, consistent with the standard laboratory RAS parameters. Group B comprised 300 zebrafish, ranging in age from 2 to 4 months post-fertilization (mpf), including 237 wild-type individuals and 63 variant-color individuals. The fish were collectively housed within the standard laboratory RAS at a stocking density of seven fish per liter.

#### 2.1.2 Zebrafish image acquisition

The imaging system employed in this study consists of a stereomicroscope coupled with a 5-megapixel PixeLINK M5D color camera. This system was utilized for all high-resolution image acquisition. The microscope and camera were connected via a standard C-mount interface and mounted on a gimbal-mounted stand, which permitted 180° rotation of the observation head and 360° rotation of the main unit. High-resolution images were captured in real time using the μEscope imaging software and subsequently saved for further analysis. Imaging was conducted within a semi-enclosed chamber. Two overhead LED strips were utilized to provide uniform, diffuse illumination, thereby minimizing reflections and shadows that could potentially compromise image quality. The images were obtained with illumination against black and blue backgrounds.

The regions of interest for individual identification encompassed the lateral body and dorsal head region (Supplementary Figure 1 A-B). Prior to imaging, zebrafish were gently transferred to a restricted-movement imaging chamber (5 × 5 × 5 cm cube, water depth 3-4 cm). Utilizing the aforementioned configuration, images were obtained of the left lateral, right lateral, and dorsal head views of each fish specimen. To ensure comprehensive coverage of body surface features, a minimum of 400 frames per fish per session were acquired. In group A, the fish were imaged over a period of 92 days, spanning from days post-fertilization 31 to 122 (Supplementary Figure 1C). The imaging frequency was adjusted during the developmental stage: daily for 1-2 mpf, every second day for 2-3 mpf, and every fourth day for 3-4 mpf. In Group B, each fish underwent a single imaging session. The video data that was obtained was systematically catalogued according to two distinct identifiers: a unique individual ID and a day age number.

### 2.2 Proposed method to identify zebrafish individuals

The zebrafish individual identification integrated framework, ESC-IDNet, constructed in this study, consists primarily of two serially connected neural networks (ESCAlignNet and IDNet), forming a cascaded model. This architecture executes integrated tasks, including target detection and standardization, feature extraction, and feature matching. The complete workflow is depicted in Figure 1. In the case of images or video frames from two temporal nodes requiring identification, the preliminary processing entails the parallel execution of ESCAlignNet, a segmentation and alignment procedure, to ensure the accurate alignment and identification of the data. Subsequently, the standardized images are registered and archived in the image database. Then, they are fed into IDNet for feature extraction, where image embeddings generate representative feature vectors. Subsequently, the feature vector sets from the two temporal nodes undergo a process of matching through the calculation of cosine distances between individual feature vectors. The determination of individual identity is based on a predefined matching threshold.

#### 2.2.1 Target detection of zebrafish

This study employed YOLOv8 for zebrafish target detection, utilizing the integrated YOLOv8-seg and YOLOv8-Pose modules for instance segmentation of the lateral body region and dorsal head region, as well as keypoint detection, respectively. In order to enhance the model’s performance in segmenting small targets, such as zebrafish, and detecting keypoints, modifications were made to YOLOv8 based on prior work [10]. Specifically, the EMA [25] attention mechanism was integrated into the Backbone and Neck, and the SPPELAN [26] and C2f-Faster [27] modules were introduced (Figure 2, Supplementary Figure 2). These modifications were designed to enhance the model’s adaptability, accuracy, and discriminative capability for subtle feature differences. The resulting improved framework, termed ESCAlignNet, is dedicated to the standardization processing of zebrafish images.

**Figure 2.**
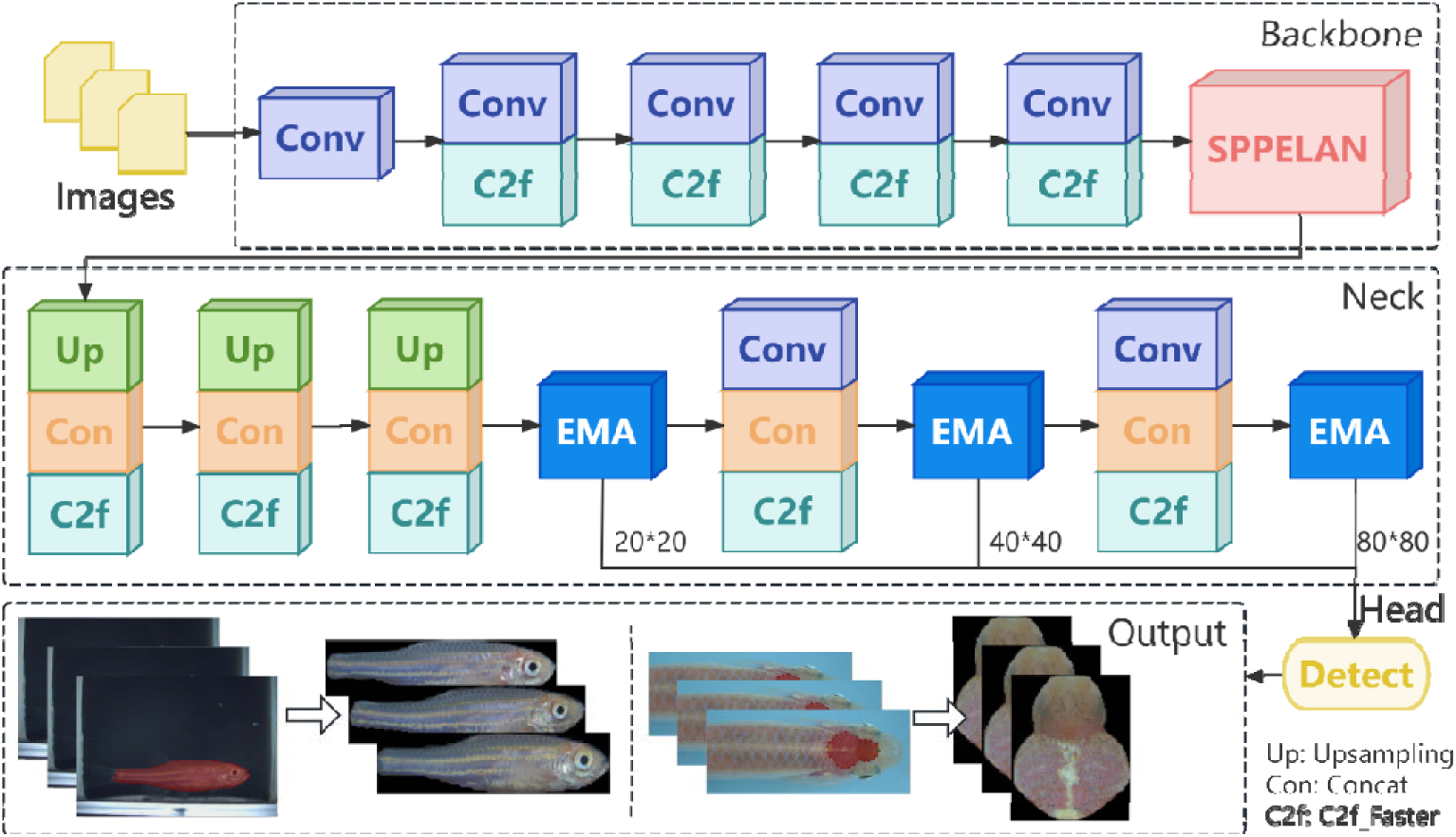
Network structure of ESCAlignNet.

Prior to the training phase, ESCAlignNet implements data augmentation operations on the dataset, encompassing Mosaic, Mixup, random perspective, and HSV augmentation techniques. The augmented images undergo a series of processing steps, beginning with a thorough extraction of features by the backbone network. Subsequently, the Neck undergoes a process of feature fusion and enhancement. In the final stage of the process, the multi-scale detection head performs precise detection and output of zebrafish. In the context of the instance segmentation task, the output delineates the boundaries of the zebrafish lateral body and dorsal head regions with a high degree of precision, achieving precise separation from the background. In the context of the keypoint detection task, the Output provides positional information regarding the lateral body and head skin, in addition to the coordinates of keypoints, including the fish eye, the midpoint of the caudal peduncle, and two dorsal points. By leveraging these keypoints, the segmented images can be aligned to a standard posture, ensuring consistency for downstream feature extraction.

With respect to hyperparameter configuration, the input image size was set to 640 × 640 pixels, with a batch size of 16 and training conducted for 300 epochs. The training program was terminated prior to its scheduled conclusion in cases where no performance enhancement was observed over the course of 50 consecutive epochs. The initial learning rate was set to 0.01, and the optimizer was configured to automatic selection (auto), with dynamic adjustments made based on the characteristics of the specific task.

The training data set was derived from a set of captured images of zebrafish, with a deliberate effort to encompass a range of developmental stages and imaging distances. This approach was taken to enhance the robustness and generalizability of the model. Annotation was performed manually using Labelme software on approximately one-sixth of the target training dataset (examples shown in Supplementary Figure 1A-B). This initially annotated set was used to train the preliminary model. Subsequently, this model was employed for the automatic annotation of new images. Subsequently, the auto-annotation results underwent manual adjustment, and the model underwent retraining with the augmented data. Through multiple iterations of this process, distinct training, validation, and test sets were established for zebrafish lateral body and head instance segmentation and keypoint detection (Supplementary Table 1). The training and validation sets were utilized for model training and parameter optimization, while the test set was reserved for performance evaluation. Prior to this, a preliminary manual screening was conducted on all images in the training, validation, and test sets to eliminate images that were blurry or distorted.

In the output layer of ESCAlignNet, a particular Laplacian kernel was integrated. This kernel performs convolution on the segmented and aligned zebrafish images to compute their corresponding variance values. The establishment of an appropriate threshold, based on this variance, enables the system to automatically filter out low-quality images, such as those that are blurry.

#### 2.2.2 Feature extraction and matching of zebrafish

After meticulously segmenting and aligning zebrafish images through ESCAlignNet, we developed the IDNet module to extract and match individual features on standardized images. The zebrafish growth curve [28] provided the foundation for the selection of images from two specific developmental windows for separate training. The initial phase (31-40 dpf; rapid-growth phase) and the subsequent phase (60-70 dpf; pre-stabilization phase of growth changes) were selected. This approach yielded distinct recognition models adapted to different growth stages. IDNet employs IResNet [29] as the primary network for zebrafish image feature extraction and incorporates a modified ArcFace loss function (Additive Angular Margin Loss, AAM Loss). The angular margin within ArcFace was adapted with the specific purpose of enhancing classification accuracy and model robustness for zebrafish individual identification (Figure 3).

**Figure 3.**
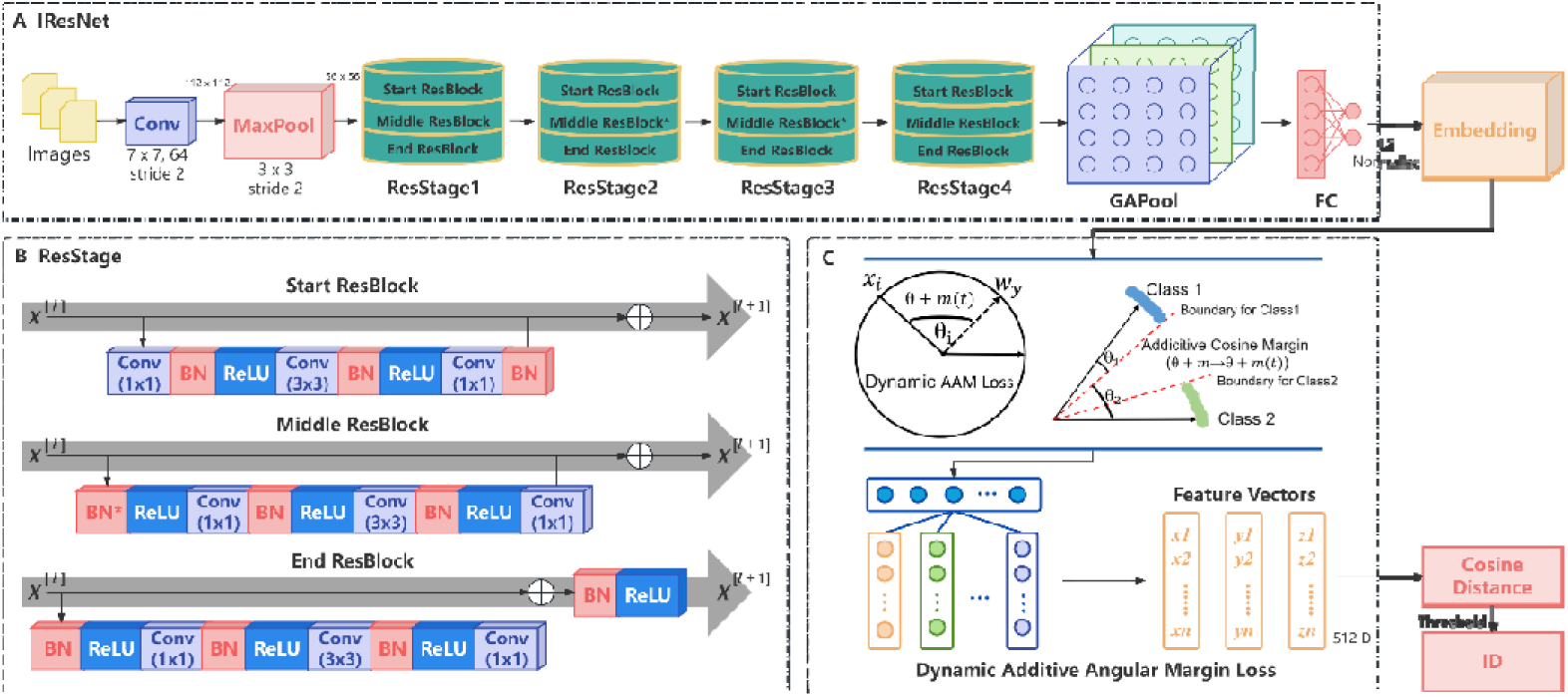
Schematic diagram of the main network architecture of IDNet. A. Main architecture of IResNet. B. Main structure of ResStage. C. Schematic diagram of dynamic additive angular margin loss optimization feature space.

The input image, standardized by ESCAlignNet, is first processed by the IResNet backbone, generating a highly discriminative feature vector. This vector undergoes L2 normalization, and the resulting embedding representation is then optimized via the AAM Loss. While the conventional ArcFace Loss presupposes static feature distributions during face recognition, it does not take into account the feature drift that is precipitated by zebrafish growth (Supplementary Figure 1C). To address this, the ArcFace loss function was modified (Supplementary Equation 1), with the angular margin ( ) being dynamically adjusted based on the zebrafish growth process. This was done to prevent feature confusion induced by developmental changes. The following presentation delineates the composition of the modified formula:

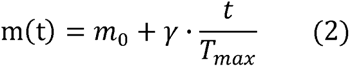

where *θ_yi_* is the angle of the correct class for sample *i*, *m(t)* is the dynamic angular margin boundary, *s* is the adjustable scale factor, *m*_0_ is the initial interval, *γ* is the growth rate coefficient, *T_max_* is the maximum duration of the experiment.

During the training phase, the initial learning rate was set to 0.01, the batch size was configured to 16, and the epochs were set to 200, employing the Stochastic Gradient Descent (SGD) optimizer. Subsequent to training, the model executes forward propagation on input zebrafish images, thereby extracting a 512-dimensional feature vector. The feature vectors in question have been demonstrated to possess both high discriminative power and robustness within the angular space. When feature matching is performed, the degree of similarity between zebrafish images is quantified by measuring the cosine distance between their feature vectors. The cosine distance is defined as follows:

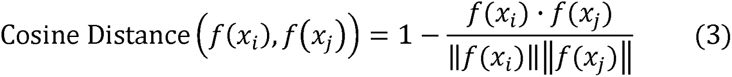

where *f*(*x_i_*) and *f*(*x_j_*) are the feature vectors of the two images. The cosine distance range is [0, 2], where 0 indicates identical vectors and 2 indicates completely dissimilar vectors.

To demonstrate the efficacy of the proposed method, ablation experiments were conducted, in which combinations of different backbone networks (Inception-ResNet v1 [30] and MobileNet [31]) with the Triplet Loss [32] (Supplementary Equation 2) were tested. These combinations employed the Euclidean distance (Supplementary Equation 3) to measure similarity. Pairs of images were assigned to the same zebrafish individual if the Euclidean distance between their feature vectors fell below a predefined threshold. For a new set of zebrafish images, the trained model extracts feature vectors, which are then matched against the feature vectors of known individuals in the database. The determination of individual identity is predicated on this matching against the established threshold.

### 2.3 Model evaluation

#### 2.3.1 Performance evaluation of instance segmentation and keypoint detection

To comprehensively evaluate the efficacy of ESCAlignNet in zebrafish image instance segmentation and keypoint detection tasks, we have identified precision, recall, mAP (mean average precision), and speed as the primary performance metrics. In order to assess the impact of segmentation and alignment operations on consistency within the same zebrafish image set, the Structural Similarity Index Measure (SSIM) [33] was employed to measure intra-class variability. SSIM is a comprehensive evaluation method that analyzes the similarity between two images in three dimensions: luminance, contrast, and structure. The value range of the similarity metric is from -1 to 1, with values closer to 1 indicating a higher degree of similarity between the two images. The calculation of the SSIM for two images, *I* and *T*, is performed as follows:

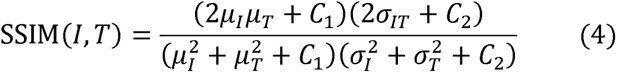

where *I* and *T* represent the local blocks (i.e., sliding window blocks) of the two images being compared. *µ_I_* and *µ_T_* are the mean luminance of *I* and *T*, respectively. 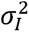 and 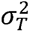 are the mean luminance of I and T, respectively. σ*_IT_* is the covariance between I and T. *c*_1_ and *c*_2_ are two constants used to stabilize the computation. The SSIM formula’s detailed specifications are provided in Supplementary Equation 4.

A comparative analysis of the SSIM differences between images of the same zebrafish that underwent segmentation and alignment processing versus unprocessed images was conducted. The experimental data were selected from zebrafish image datasets spanning 31-122 dpf. A total of three subjects were selected at random, and 300 images were randomly sampled from each subject. For each individual, the mean and standard deviation of the SSIM were calculated for all possible image pairs, both before and after segmentation and alignment processing. Subsequently, a comparative analysis was conducted on the alterations in these values before and after processing to assess intra-class variability.

#### 2.3.2 Performance Evaluation of Feature Extraction and Matching

During the construction of the test dataset, clear zebrafish images from various developmental stages that were not involved in the training process were selected from the image database. These images were formatted according to the Labeled Faces in the Wild (LFW) standard to create a validation dataset containing “positive sample” pairs and “negative sample” pairs. In “positive sample” pairs, both images originate from the same zebrafish individual, while “negative sample” pairs consist of images from different individuals. For this study, each LFW-formatted test sample was subjected to 10 independent random sampling iterations, with each iteration comprising 300 positive pairs and 300 negative pairs. Following the aggregation of all sampling results, six distinct LFW-formatted files were established, each containing 3,000 positive sample pairs and 3,000 negative sample pairs. ROC (Receiver Operating Characteristic) curves were subsequently plotted to determine the optimal decision threshold during evaluation testing.

ROC curve analysis was performed by evaluating the False Positive Rate (FPR) and True Positive Rate (TPR) across varying thresholds. The threshold ranges for both the Euclidean distance and the cosine distance were set to span from 0 to 2, with an increment step size of 0.01. The model’s performance was quantified using the Area Under the ROC Curve (AUC). An AUC value approaching 1 signifies enhanced classification efficacy. The selection of the optimal threshold point was based on the variations in the Area Under the Curve (AUC) value. The model’s accuracy and validation rate were subsequently calculated.

Through the implementation of a comprehensive testing process, which involved the combination of diverse backbone networks and loss functions, the optimal combination was identified, exhibiting both superior accuracy and expedited processing speed. This optimal model was then employed to recognize longitudinally captured zebrafish images to assess its long-term stability. Concurrently, the matching accuracy between zebrafish images from different growth stages and images registered in the database was compared. The performance alterations in the model for long-term image recognition tasks were evaluated by recording recognition accuracy at varying time intervals, followed by quantitative analysis.

## 3. Results

### 3.1 Construction of zebrafish growth images database

For the longitudinally imaged Group A (150 zebrafish), a total of 13,900 lateral body videos were acquired, with each video comprising 200 frames, alongside 56,400 dorsal view images. For the short-term imaged Group B (300 zebrafish), 452 videos were obtained, including 300 lateral body videos and 152 dorsal view videos. Subsequent to the extraction of frames from all videos and the elimination of distorted or incomplete fish body images, approximately 300,000 frames were retained. The ESCAlignNet described in Section 2.2.1 underwent multiple iterations. The final model, applied to instance segmentation and keypoint recognition, was trained and tested on datasets comprising 15,642 lateral body images and 8,047 dorsal head images, respectively (Figure 4A, Supplementary Table 1).

**Figure 4.**
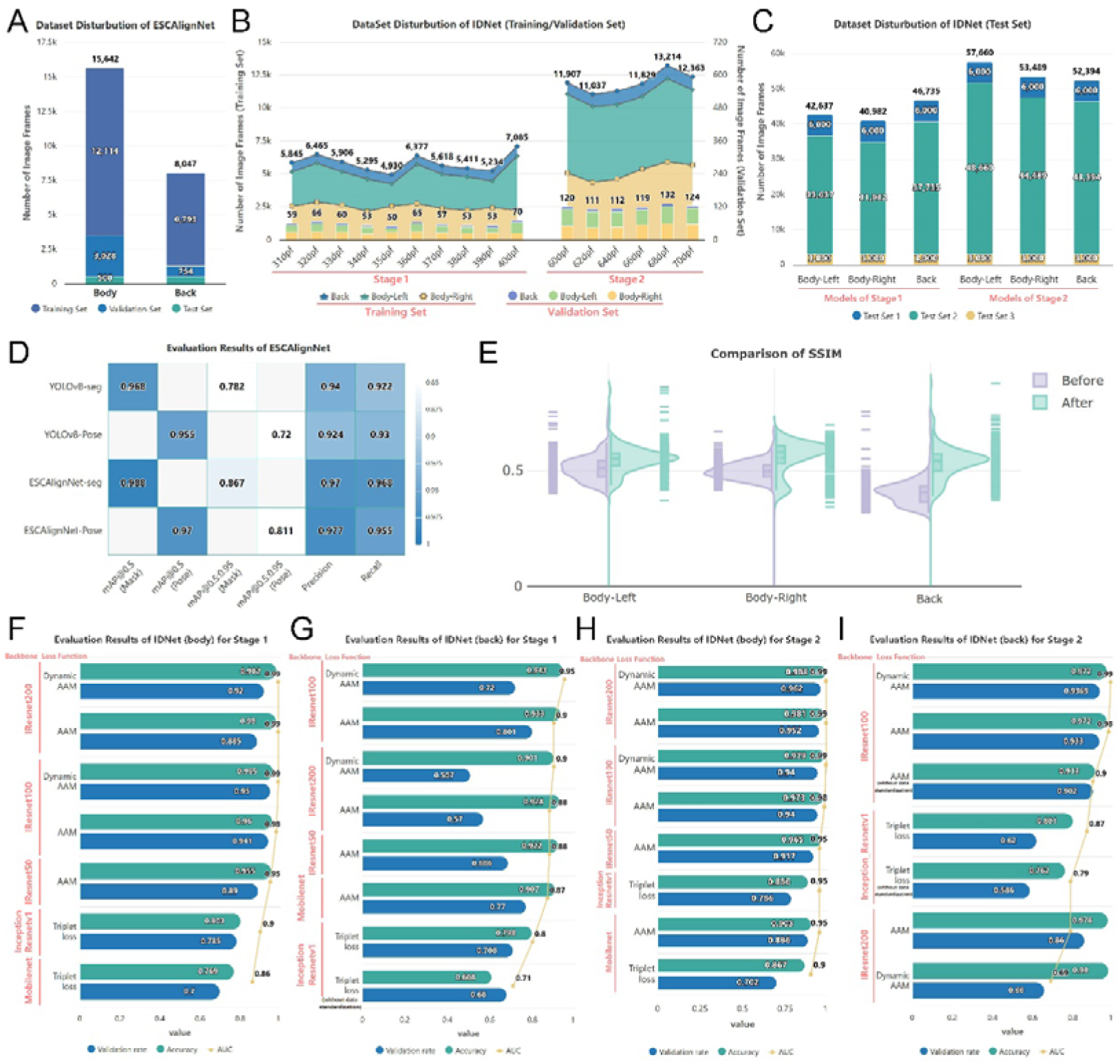
Data distribution for training and testing the ESC-IDNet framework during 31-122 dpf and corresponding evaluation results. The lateral body and dorsal head are designated as “body” and “back”, respectively, in Figure. A. Training, validation, and test sets for ESCAlignNet training on zebrafish lateral body (body) and dorsal head (back) instance segmentation and keypoint recognition. B. IDNet module training/validation datasets (line graph/bar chart), with daily sampling in Stage 1 (31-40 dpf) and alternate-day sampling in Stage 2 (60-70 dpf). C. Test set data for IDNet models during Stage 1 and Stage 2: number of test images categorized by imaging view (left lateral, right lateral, dorsal). D. Training and testing evaluation results for YOLOv8 versus ESCAlignNet. E. Distribution comparison of SSIM values before and after standardization for images of three individuals across body regions. F. Test Set 1 evaluation results for the Stage 1 IDNet (Body) model. G. Test Set 1 evaluation results for the Stage 1 IDNet (Back) model. H. Test Set 1 evaluation results for the Stage 2 IDNet (Body) model. I. Test Set 1 evaluation results for the Stage 2 IDNet (Back) model.

The zebrafish image collection was processed through a series of automated procedures using a model trained via ESCAlignNet. These procedures included segmentation, alignment standardization, and automated filtering. Standardized high-quality images were systematically archived by date and unique identification number, culminating in a comprehensive dataset encompassing diverse individual characteristics and developmental stages. The final dataset comprises approximately 200,000 clear lateral body images and 90,000 clear dorsal head images, with each image possessing a unique identifier, timestamp, and corresponding developmental stage designation.

The training and validation data for IDNet were exclusively sourced from the longitudinally imaged Group A dataset, comprising 150 zebrafish. Meanwhile, test data from external fish populations were derived from the Group B dataset, which contains 300 zebrafish. The training and validation sets were partitioned into two distinct stages: Stage 1 (31-40 dpf) and Stage 2 (60-70 dpf) (Figure 4B). Subsequent to the development of the stage-specific IDNet models, three distinct test sets were evaluated (Figure 4C). Test Set 1 employed the generated LFW-formatted files to assess both stage-specific models. Test Set 2 administered chronological individual identification tests utilizing the Group A images. Both models were evaluated at every imaging timepoint between 31-122 dpf, with approximately ten images per individual per timepoint. Test Set 3 evaluated both models using Group B data, comprising 300 zebrafish, with approximately ten randomly selected images per fish. Supplementary Table 2 provides a more detailed specification of the training and testing data.

### 3.2 Results of segmentation and alignment

The ESCAlignNet module was utilized to achieve 0.970 and 0.977 for precision in the test dataset for instance segmentation and keypoint recognition, respectively. These values represent increases of 0.030 and 0.053 compared to the baseline YOLOv8 model (Figure 4D). Concurrently, enhancements were observed in mAP and Recall, though the model layer count increased from 261 to 377, and inference speed decreased from 240 frames per second (FPS) to approximately 108 FPS (Supplementary Table 3).

Following the segmentation and alignment of 900 images (3 × 3 × 100) randomly selected from three individuals, the SSIM values for all body regions per individual demonstrated significant increases (*p* < 0.01). The mean SSIM for left lateral, right lateral, and dorsal head images increased from 0.5138, 0.5016, and 0.4075 to 0.5726, 0.5523, and 0.5576, respectively. This finding suggests that standardization processing significantly enhances intra-class consistency within individual samples (Figure 4E, Supplementary Table 4).

### 3.3 Results of feature extraction and matching

Utilizing the LFW-formatted Test Set 1, a comparative evaluation was conducted on stage-specific IDNet models, with an analysis of performance across various combinations of backbone networks and loss functions (Figure 4F-I, Supplementary Tables 5-8). The experimental results demonstrated that IDNet (Body) models for Stages 1 and 2, trained with IResNet200 and Dynamic AAM loss, achieved optimal recognition accuracy on lateral body test data (Figure 4F, 4H). In contrast, IDNet (Back) models for both stages, trained with IResNet100 and Dynamic AAM loss, exhibited higher accuracy on dorsal head test sets of equivalent size (Figure 4G, 4I). A comparison of the performance metrics between the two model types revealed no statistically significant differences in Area Under the Curve (AUC) or Validation Rate across the various stages of the study. However, IResNet200 led to a substantial increase in model weight size and a significant reduction in FPS. Therefore, following a thorough evaluation of metrics including accuracy, validation rate, AUC, and FPS, IResNet100 was selected as the backbone network integrated with dynamic AAM loss to form the core module of IDNet. The optimal identity discrimination threshold was determined to be 1.00 (Supplemental Tables 5-8).

In order to assess the framework’s long-term stability, Test Set 2 was employed to evaluate the sustained recognition capability of ESC-IDNet models trained on different body regions across both stages (Figure 5).

**Figure 5.**
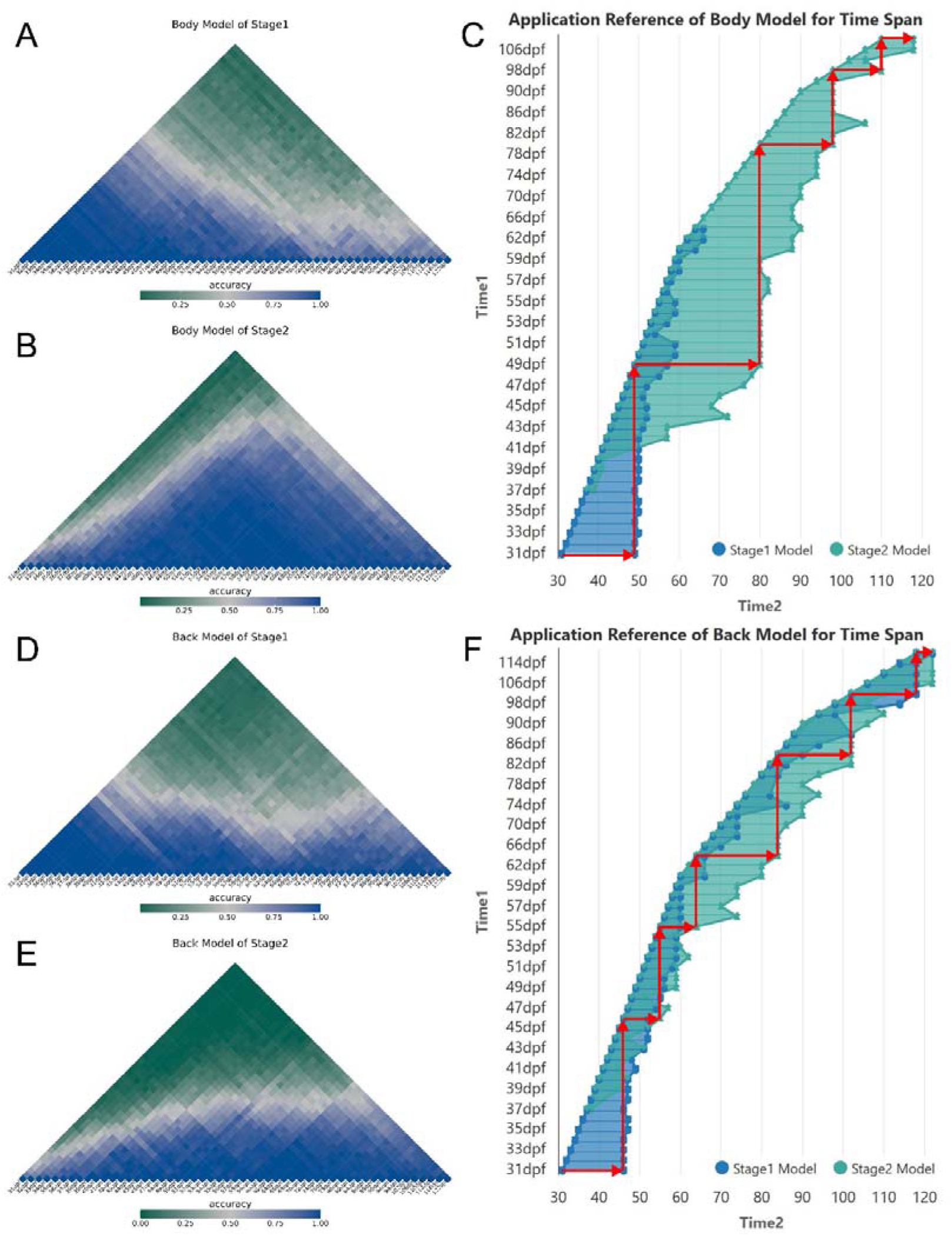
Long-term performance testing of zebrafish individual identification using Test Set 2. A-B. Recognition accuracy over time for models trained on Stage 1 and Stage 2 lateral body data. C. Time span maintaining ≥95% accuracy with update strategies implemented at critical timepoints for the lateral body model. D-E. Recognition accuracy over time for models trained on Stage 1 and Stage 2 dorsal head images. F. Time span maintaining ≥95% accuracy with update strategies implemented at critical timepoints for the dorsal head model. Red arrows indicate an optimized strategy for model and database update timing.

An analysis of the results from Test Set 2 indicated a significant correlation between the pattern of recognition accuracy decay over time and model selection, as well as the temporal reference point of the baseline images. It was observed that all models demonstrated a decline in recognition accuracy as the time interval between reference and query images increased. However, when reference images were updated at 1-2 day intervals, recognition accuracy remained consistently at 100% (Figures 5A, 5B, 5D, 5E). Overall, the models exhibited stage-dependent robustness to subsequent developmental data. Specifically, models that were trained on Stage 2 data (Figures 5B and 5E) demonstrated high accuracy over significantly longer durations than those trained on Stage 1 data (Figures 5A, 5D). Furthermore, when subjected to a specified accuracy threshold (e.g., Accuracy ≥ 0.95), distinct inflection points were identifiable beyond which recognition accuracy declined below the required threshold. The red arrows denote the critical junctures that serve as reference points for database updates (Figure 5C, 5F). Updating both the database (by replacing original reference images with those from a given time point) and the recognition model is essential to restore and maintain high-accuracy identification performance. For instance, commencing identification at 31 dpf using the Stage 1 lateral body model (Figure 5C), the initial database update and model switch to Stage 2 is required at approximately 49 dpf to maintain ≥95% accuracy, with subsequent accuracy sustained until 80 dpf necessitating another update. Furthermore, maintaining the same level of accuracy necessitated a smaller number of database updates for lateral body recognition (Figure 5C, 4 updates) than for dorsal head recognition (Figure 5F, 6 updates). It is noteworthy that during Stage 1, the dorsal head model (illustrated in blue in Figure 5F) demonstrated a longer effective recognition span (31-122 dpf) in comparison to the lateral body model (31-64 dpf).

In Test Set 3 evaluations, all stage-specific and region-specific recognition models achieved 100% accuracy when identifying the 300 individuals from the external population.

### 3.4 Individual identification online platform

The framework described in this study was implemented on a Dell T3650 workstation that was configured with the following hardware specifications: an Intel i7-11700 processor (8 cores, base frequency 2.5 GHz, maximum turbo frequency 4.9 GHz), 32 GB of RAM, and an NVIDIA RTX A4000 graphics card. The software dependencies included PyTorch version 1.12.0 and CUDA version 11.3. The ESC-IDNet framework, utilized in this study for the purpose of real-time zebrafish individual identification, can be locally deployed. In order to validate the efficacy and practical utility of the proposed methodology, a non-real-time online identification platform was developed. This platform was designated FishIndivID (http://bioinfo.ihb.ac.cn/fishindivid) and was developed based on the Django web framework. The development of this platform was motivated by the necessity to facilitate the identification and matching of individuals across images captured at different time points. The primary focus of the platform is on the lateral and dorsal head regions of zebrafish. It is possible for users to upload raw images obtained at two separate time points in order to obtain individual matching results. As illustrated in Figure 6, the operational process of the platform is composed of three fundamental steps. Initially, lateral-view images captured at Time Point 1 are uploaded. Then, the corresponding images captured at Time Point 2 are uploaded. Subsequently, a matching request is submitted, and the platform returns the identification results, which include a cosine distance matrix (visualized as a heatmap) representing pairwise similarities between individuals across the two time points, and the corresponding matching results inferred from the cosine distances, presented as a Sankey diagram.

**Figure 6.**
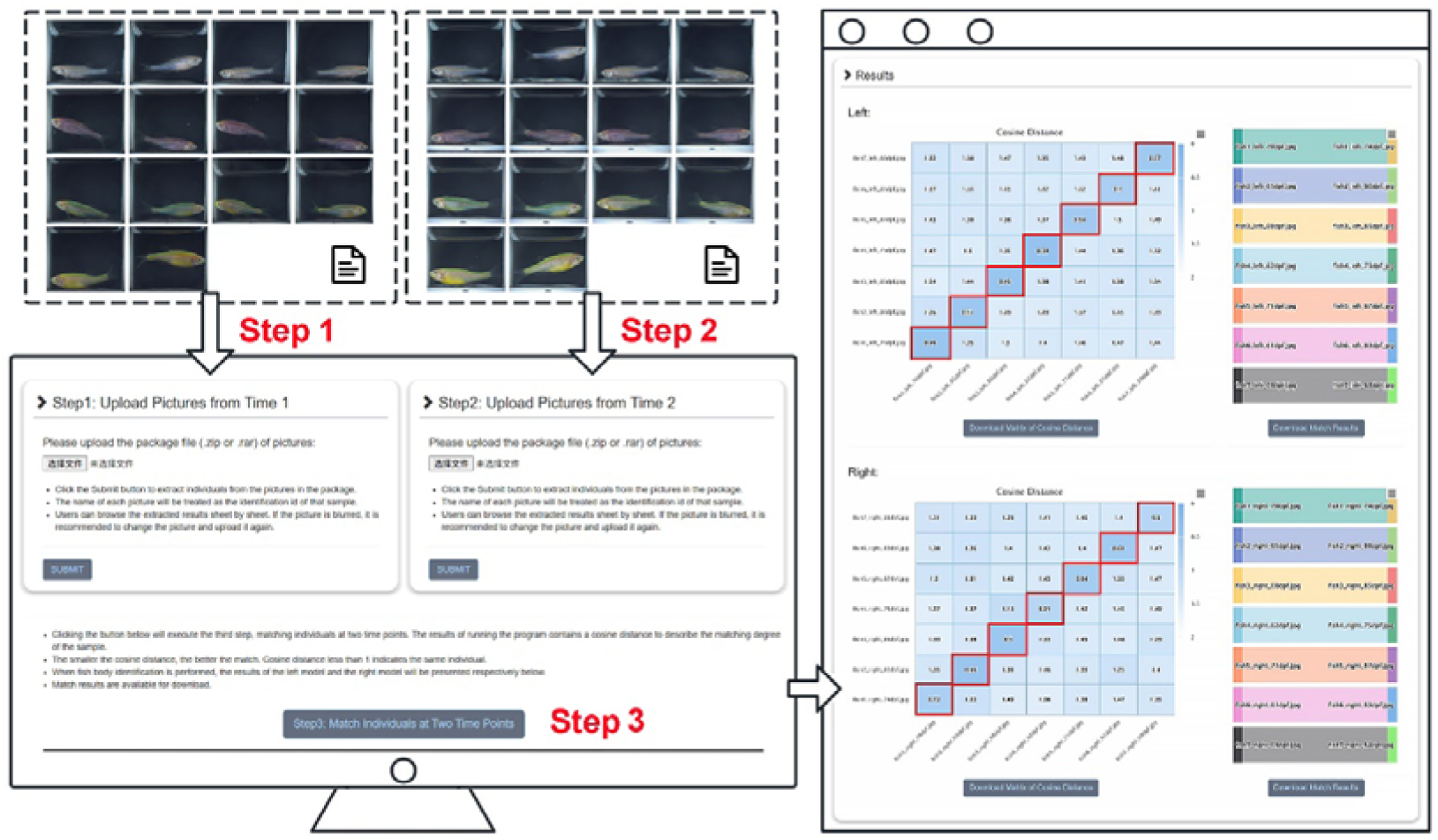
The main steps involved in individual identification on an online platform and the output results.

## 4. Discussion

The advent of deep learning technologies has precipitated a paradigm shift in methodologies employed for biological individual identification. However, significant challenges remain for fish species due to large populations and minimal inter-individual morphological variation. This is particularly acute in small fish like zebrafish, where rapid growth cycles and dynamic phenotypic changes further hinder individual recognition. Our study systematically addressed these challenges through the development of the ESC-IDNet framework, which integrates targeted region selection, advanced image standardization, optimized feature extraction, and a dynamic update strategy to achieve robust, long-term individual identification.

First, feature selection and data scale critically impact recognition performance. While previous zebrafish studies predominantly used the lateral body for its distinct texture and large area [22, 23], this region suffers from bending, occlusion, and illumination variance. In contrast, the dorsal head exhibits a distinctive shape with fine speckling (Supplementary Figure 1B) and clear boundaries, facilitating stable acquisition. This study pioneers the dorsal head as a feature region for identification. Large-scale datasets are essential for CNN generalization, yet such data remain scarce for non-human identification [34]. To address this, we developed a recirculating aquaculture system with integrated imaging, enabling longitudinal data collection across developmental stages. This temporal dataset enhances model adaptability beyond transient features. Evaluation on Test Set 2 showed the lateral body model achieved high accuracy (≥95%) over extended periods, while the dorsal head model exhibited an augmented model validity period (the duration of Stage 1 and Stage 2 models). Biologically, this difference stems from the greater topological stability of the cranial structure versus the lateral body. The latter undergoes pronounced feature drift during juvenile-to-adult transition, causing the Stage 1 lateral model to fail beyond 64 dpf (Accuracy < 95% across all temporal intervals). Although the dorsal head model maintained high long-term accuracy, its smaller detection area increases acquisition difficulty, constraining the training dataset to ∼20% of the lateral body data (Figure 4B). This compromises model robustness and results in lower absolute accuracy than the body model at equivalent time points. Addressing this challenge requires specialized image acquisition apparatuses.

Second, image standardization significantly reduces intraclass variability. Similar to the image standardization achieved through detection and alignment in facial recognition, zebrafish individual identification also necessitates the reduction of intra-class variability induced by factors such as posture, illumination, and background. However, achieving this standardization underwater presents greater challenges due to dynamic light fluctuations (from bubbles, suspended particles, water flow) and the elusive morphology of dorsal, pelvic, and caudal fins during movement, hindering the acquisition of stable, standardized images (Supplementary Figure 1A). Puchalla et al. [23] demonstrated that Convolutional Neural Networks primarily rely on zebrafish stripe patterns for feature discrimination rather than on head, tail, or fin morphology. Therefore, this study employed instance segmentation technology during object detection, meticulously segmenting the pelvic, dorsal, and caudal fins to enable precise isolation of the lateral body and dorsal head regions. This approach offers advantages over bounding boxes by providing refined morphology, eliminating complex background interference, retaining only fish-related pixels, enhancing feature extraction accuracy, and consequently improving recognition performance. After undergoing keypoint detection and alignment, the segmented regions proceed to feature extraction. In order to achieve the above automatic standardization process, we modified YOLOv8 to create the ESCAlignNet module within the ESC-IDNet framework. Key modifications include integrating an EMA mechanism into the backbone to preserve precise ROI spatial information and facilitate positional feature extraction for accurate detection [25] (Supplementary Figure 2D); replacing the original SPPF module with SPPELAN at the backbone base to enhance detection accuracy through multi-scale feature extraction and efficient layer aggregation while significantly boosting speed for real-time applications, particularly beneficial for fast-moving small targets (Supplementary Figure 2A); and forming a C2f-faster structure by replacing the Bottleneck in C2f with a FasterNet Block to mitigate increased parameter/complexity, reducing both while maintaining an adequate receptive field and nonlinear representation capacity (Supplementary Figure 2B, C). Although these changes slightly increase model intricacy and reduce recognition speed, performance remains sufficient for detection. Experimental results demonstrate that ESCAlignNet enhances instance segmentation and keypoint detection performance (Figure 4D, Supplementary Table 3) and effectively reduces intra-individual image variability (Figure 4E, Supplementary Table 4), significantly increasing intra-class similarity. This precise region extraction and alignment normalizes input data, minimizing the impact of capture angles and posture while substantially reducing background noise and image complexity. Consequently, this approach not only improves zebrafish recognition rates across diverse imaging environments but also establishes a robust foundation for subsequent recognition tasks.

Third, optimizing the recognition backbone and loss function maximized discrimination of subtle phenotypic features. The field of facial recognition has seen the development of numerous mature feature extraction and matching algorithms, including DeepFace [35], FaceNet [36], ArcFace [32], and OpenFace[37]. Among these, FaceNet and ArcFace represent two classical architectures. FaceNet constructs a high-dimensional feature space using triplet loss, ensuring that similar samples are proximate while dissimilar samples are distantly separated. In contrast, ArcFace incorporates an angular margin mechanism to optimize the distribution of feature embeddings, thereby enhancing the model’s discriminative power for inter-class differences. A comparison of Inception-ResNet and MobileNet with FaceNet’s original backbone networks (ZFNet [38] and GoogLeNet V1 [39]) reveals that the former offer greater architectural flexibility, deeper layers, and superior capability for extracting complex features. In the context of backbone network selection based on triplet loss, the study found that Inception-ResNet v1 significantly enhanced recognition accuracy, particularly in the classification of similar individuals. This enhancement can be attributed to the integrated Inception modules and residual connections within the network, which facilitate optimal gradient propagation. While MobileNet, as a lightweight network, achieved higher FPS, its lower accuracy rendered it suboptimal for fine-grained classification tasks. Consequently, in the context of zebrafish recognition, Inception-ResNet v1 demonstrated superior accuracy, while MobileNet exhibited higher FPS (Figure 4F-I, Supplementary Tables 5-8). In both facial recognition and fine-grained classification tasks, ArcFace generally outperforms FaceNet across multiple benchmarks [32]. This superiority extends to the realm of zebrafish individual identification. On the one hand, FaceNet’s triplet loss demonstrates sensitivity to sample mining strategies and exhibits a paucity of explicit constraints for intra-class compactness, rendering it susceptible to noise in scenarios characterized by minimal intra-class variation, as is the case in zebrafish. ArcFace loss, by optimizing angular margin between feature vectors, accentuates inter-individual differences among zebrafish. The dynamic adjustment of the angular margin serves to mitigate feature confusion induced by growth-related changes. On the other hand, ArcFace loss necessitates the conversion of image features into 512-dimensional vectors, in contrast to the 128-dimensional vectors employed by triplet loss. This increased dimensionality facilitates superior feature representation capacity and information retention. Consequently, dynamic ArcFace loss emerges as a preeminent approach for zebrafish individual identification. This study employed an enhanced version of ResNets, designated as IResNets. IResNets enhance ResNet through modifications to three key components: information flow, residual sub-modules (ResBlocks), and projection shortcuts. These refinements address challenges related to optimization and information degradation in deep network training, thereby enabling IResNets to surpass ResNet in terms of both accuracy and convergence [29]. As the depth of the network increased across the three backbone networks (IResNet50/100/200) utilized with ArcFace loss, the feature extraction capability and recognition accuracy improved progressively, albeit with escalating computational resource demands. Based on a comprehensive evaluation of multiple backbone network and loss function combinations (Figure 4F-I, Supplementary Table 5-8), IResNet100 was ultimately selected as the backbone network for IDNet, paired with dynamic ArcFace Loss as the loss function. Integrating this with the ESCAlignNet module, the proposed methodology is designated the ESC-IDNet framework for integrated zebrafish individual identification.

Fourth, feature update mechanisms across developmental stages facilitate long-term individual recognition. The continuous alterations in the morphology of the zebrafish surface that occur throughout its developmental stages render static matching strategies inadequate for sustaining long-term stable identification. During the course of the long-term stability testing (Test Set 2) of the zebrafish individual identification in this study, models that were trained on Stage 2 data demonstrated significantly higher stability and recognition rates for both the lateral body and dorsal head regions in comparison to Stage 1 models (Figure 5). This enhancement can be attributed to the training of Stage 2 models on data obtained during the period closer to the morphologically more stable adult stage. Consequently, the features learned exhibit greater persistence during subsequent development, resulting in slower decay and an extended time window for effective recognition. Conversely, Stage 1 models capture rapidly changing larval-stage features, whose representativeness diminishes markedly in later developmental phases, accelerating the decline in recognition accuracy. This finding underscores the critical importance of training data collection timing for longitudinal applications. Subsequent training of models on later, morphologically stabilized developmental stages results in models that exhibit enhanced robustness for adult individuals. This evidence further corroborates the assertion that training models exclusively on data from a solitary time point is inadequate for achieving long-term tracking. To address this challenge, this study proposes and implements a dynamic update strategy. Guided by data-driven analysis (Figure 5C, F), this strategy provides a guidance for updating model and databases. Specifically, the updating process is initiated at inflection points in accuracy trends, where outdated reference images are systematically replaced with images from the current time point. This process effectively re-anchors the model to the current state, thereby mitigating feature drift induced by growth. Moreover, the update strategy employed differs fundamentally from conventional approaches, including batch retraining methods and methods relying solely on sparse, fixed-time-point reference images. The system has been engineered to strike a balance between two competing imperatives: the need for accurate recognition and the requirement for operational feasibility. The utilization of accuracy monitoring to guide the timing of model and database updates enables continuous tracking. This approach circumvents the encumbrance of incessant manual validation and the onerous demands of extensive retraining, thereby markedly enhancing the efficacy of experimental procedures.

The robust individual identification framework established in this study, capable of tracking zebrafish from larval to adult stages with minimal manual intervention, fundamentally transforms the scale and granularity of longitudinal experimental design. ESC-IDNet transcends mere technical achievement by enabling previously impractical research paradigms. It allows for the precise investigation of critical biological processes such as the ontogeny of complex behaviors, the longitudinal progression of disease phenotypes within genetically defined individuals, and the nuanced responses to chronic drug or toxicant exposures at an individual level. This capability moves the field beyond population-level averages, permitting the dissection of individual variability—a crucial factor in understanding differential susceptibility, resilience, and treatment outcomes. Our approach thus provides a powerful tool not only for basic biological discovery but also for pre-clinical research, where tracking the lifelong trajectory of each subject is paramount.

Finally, while the methodology proposed in this study demonstrates strong performance, it faces several challenges in practical applications. The image acquisition process entails considerable hardware requirements, and the ESC-IDNet framework itself demands substantial computational resources. Moreover, as the zebrafish image database grows with the inclusion of additional registered individuals, the time required for individual identification increases, raising hardware requirements for real-time monitoring applications. The inflection points identified for the dynamic update strategy are derived from test data specific to the experimental cohort studied. The presence of variability in individual growth rates may also introduce minor influences on the determination of the optimal update timing for specific populations. To enhance its practical utility, future improvements and extensions will focus on the following areas:

1. Model Optimization: Refining the model architecture to reduce computational complexity. Future work will explore the use of more advanced networks, such as Vision Transformers [40].
2. Multimodal Data Integration: Incorporating additional data types, such as zebrafish movement behaviors, to establish a multimodal individual identification system, enhancing recognition accuracy and reliability.
3. Multi-Stage Modeling and Testing: Dividing the growth curve into more distinct developmental stages for modeling, integrating more robust features to extend the effective recognition time window. Analysis of the most temporally stable local stages, combined with omics data, will facilitate the identification of molecular regulatory mechanisms most strongly associated with dermal feature changes.
4. Cross-Species Application: Extending the methodology to enable individual identification in other fish species, broadening its applicability. In summary, the significance of zebrafish individual identification lies in its applications across developmental, behavioral, and genetic research, enabling researchers to track individual differences and gain deeper insights into biological processes. Although the method presented here was developed and validated for zebrafish identification, it constitutes an adaptable framework. By adjusting image acquisition devices and model parameters, this approach can be extended to individual identification in other fish species, thereby supporting research in areas such as genetic breeding of aquacultured species, ecological monitoring, population dynamics analysis, and studies of environmental change impacts.

## 5. Conclusions

In addressing the practical need for individual identification of zebrafish in scientific research, this study proposes a computer vision-based deep learning framework, ESC-IDNet. This framework has demonstrated remarkable efficacy in zebrafish individual recognition tasks. The establishment of an online demonstration platform, FishIndivID, was undertaken to facilitate the potential application of this technology in developmental, behavioral, and genetic studies of zebrafish. Through large-scale image acquisition and processing across growth stages, model enhancement (integrating the two-stage deep learning networks ESCAlignNet for image standardization and IDNet for identification), and a dynamic updating strategy, this work systematically addresses the challenges of automatic image standardization and cross-temporal recognition. The experimental results demonstrate that the trained model derived from this approach achieves reliable identification of zebrafish individual identities. Future work will focus on enhancing data diversity, optimizing the model architecture, and exploring multimodal data integration to further improve recognition accuracy and adaptability. Furthermore, the proposed methodology offers a scalable solution for individual identification in other fish species by expanding datasets and model fine-tuning.

## Ethics statement

All experiments and animal treatments were carried out according to the principles of Animal Care and Use Committee of Institute of Hydrobiology, Chinese Academy of Sciences.

## CRediT authorship contribution statement

**Danying Cao**: Conceptualization, Methodology, Software, Formal analysis, Investigation, Writing - original draft. **Cheng Guo**: Formal analysis, Investigation, Software, Visualization. **Yingyin Cheng**: Validation, Resources, Supervision. **Wanting Zhang**: Data curation. **Mijuan Shi**: Conceptualization, Investigation, Writing - review & editing, Funding acquisition. **Xiao-Qin Xia**: Conceptualization, Investigation, Writing - review & editing, Project administration, Funding acquisition.

## Acknowledgments

This work was supported by the National Key R&D Program of China (Grant No. 2021YFD1200804 and 2023YFD2401603), and the National Natural Science Foundation of China (Grant No. 32473146).

## Data Availability

Data will be made available on request.

