## Supplemental Materials for "Longitudinal Identification of Zebrafish Individuals by Deep Learning"

### Supplementary Figures

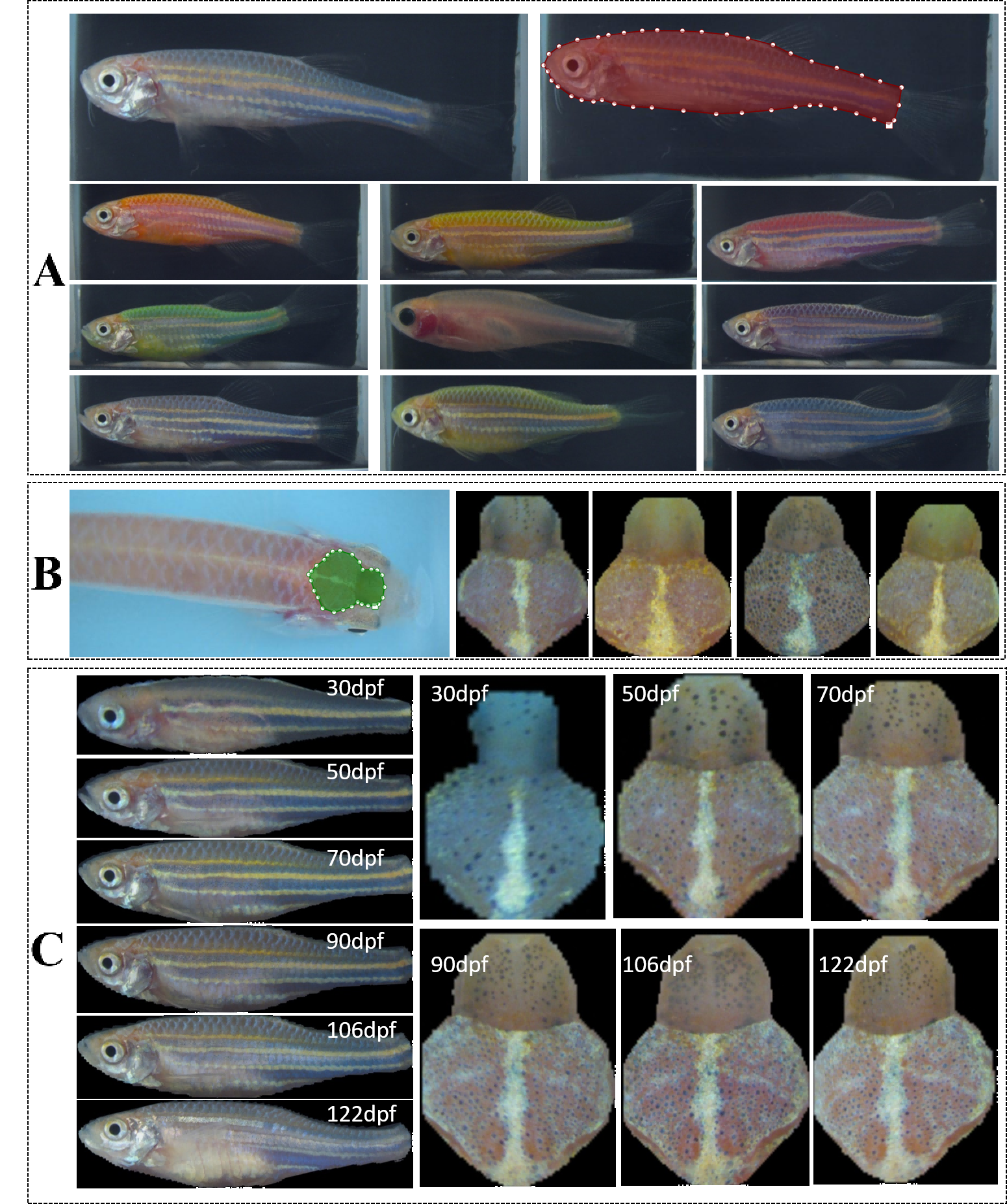

Supplementary Figure 1. Zebrafish example images. A. Selection of zebrafish lateral head region and example images of different zebrafish strains. B. Selection and example of zebrafish dorsal region. C. Dual-region images of the same zebrafish from 31 to 122 dpf.

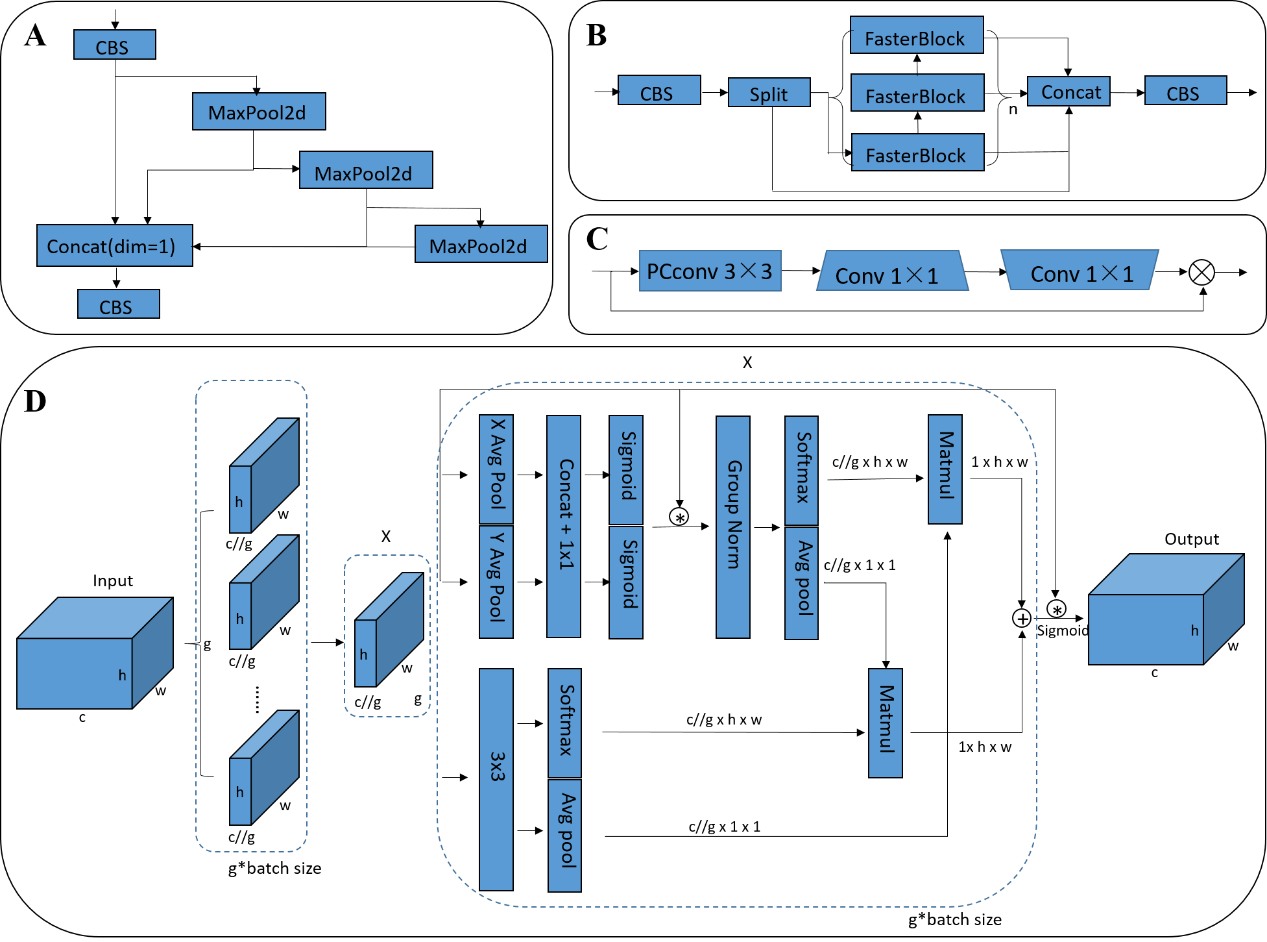

Supplementary Figure 2. Schematic diagram of the improved modules of the ESCAlignNet network structure. A. SPPELAN module. B. C2f_Faster module. C. Faster Block module. D. EMA attention mechanism module.

### Supplementary Equations

Supplementary Equation 1:

ArcFace is a loss function used for facial recognition, fully known as Additive Angular Margin Loss (AAM loss). It enhances the discriminative ability of features by increasing the interval in the angular space. The loss function is defined as:

$$L=-\frac{1}{N}\sum_{i=1}^{N} log\frac{e^{s\left( \cos\left( \theta_{y_{i}}+m \right) \right)}}{e^{s\left( \cos\left( \theta_{y_{i}}+m \right) \right)}+\sum_{j\neq y_{i}} e^{cos\theta_{j}}} (1)$$

Where $s$ is an adjustable scale factor, $\theta_{y_{i}}$ is the angle of the correct class for sample $i$, and $m$ is the angle margin. The margin $m$ forces$\theta_{y_{i}}$ of same-class samples to become smaller, making features more compact on the hypersphere. The angle between different classes increases by at least $m$, improving inter-class separability.

Supplementary Equation 2:

Triplet Loss is another commonly used loss function in face recognition. Its core component, the "triplet," consists of three samples. It enhances class discriminability by optimizing distance relationships in the feature embedding space. The loss function is defined as:

$$T= max(0,\left\| f\left( x_{a} \right)-f(x_{p}) \right\|_{2}^{2}-\left\| f\left( x_{a} \right)-f\left( x_{n} \right) \right\|_{2}^{2}+\alpha) (2)$$

Where $x_{a}$ is the anchor sample (reference image), $x_{p}$ is another image of the same identity as the anchor (positive), $x_{n}$ is an image from a different identity (negative), $\alpha$ is the margin hyperparameter (fixed margin), $f\left( \cdot\right)$ denotes a 128-dimensional embedding vector extracted by the backbone network.

Supplementary Equation 3:

The Euclidean distance formula is:

$$\mathrm{Distance}\left( f\left( x1 \right),f(x2) \right)=\left\| f\left( x1 \right)-f(x2) \right\|_{2}=\sqrt{\sum_{i=1}^{n} {({f\left( x_{1} \right)}_{i}-{f\left( x_{2} \right)}_{i})}^{2}} (3)$$

Where $f\left( x1 \right)$ and $f(x2)$ are feature vectors of images, $n$is the feature dimension (128−dimensional when using Triplet Loss), and $\left\| \cdot\right\|_{2}$ denotes the L2-norm (Euclidean distance). After L2-normalization, the Euclidean distance between two feature vectors ranges within [0, 2].

Supplementary Equation 4:

SSIM (Structure Similarity Index Measure) is an objective metric for evaluating structural similarity between two images. Its core principle calculates similarity by comparing three dimensions: luminance (*l*), contrast (*c*), and structural information (*s*). The formula is:

$$\mathrm{SSIM}\left( x, y \right)={[l(x,y)]}^{\alpha}\cdot{[c(x,y)]}^{\beta}\cdot{[s(x,y)]}^{\gamma} (4)$$

$$l\left( x,y \right)= \frac{2\mu_{x}\mu_{y}+C_{1}}{\mu_{x}^{2}+\mu_{y}^{2}+C_{1}} , C_{1}={(K_{1}L)}^{2} (5)$$

$$c\left( x,y \right)= \frac{2\sigma_{x}\sigma_{y}+C_{2}}{\sigma_{x}^{2}+\sigma_{y}^{2}+C_{2}} , C_{2}={(K_{2}L)}^{2} (6)$$

$$s\left( x,y \right)= \frac{\sigma_{xy}+C_{3}}{\sigma_{x}\sigma_{y}+C_{3}} , C_{3}=C_{2}/2 (7)$$

Where $\mu_{x}, \mu_{y}$ are local window means, $L$ is the dynamic range of pixel values, $\sigma_{x}{,\sigma}_{y}$ are local standard deviations, and $\sigma_{xy}$ is the covariance. $K_{1}=0.01$ and $K_{2}=0.03$. Setting $\alpha=\beta=\gamma=1$, the SSIM formula simplifies to:

$$\mathrm{SSIM}\left( x,y \right)=\frac{\left( 2\mu_{x}\mu_{y}+C_{1} \right)\left( 2\sigma_{xy}+C_{2} \right)}{\left( \mu_{x}^{2}+\mu_{y}^{2}+C_{1} \right)\left( \sigma_{x}^{2}+\sigma_{y}^{2}+C_{2} \right)} (8)$$

### Supplementary Tables:

Supplementary Table 1 provides the detailed distribution statistics of the data used to train, validate, and test the zebrafish instance segmentation and alignment model (ESCAlignNet). The dataset includes 450 fish and covers all time points (31–122 dpf) in the age distribution of captured images.

Supplementary table 1. Dataset sizes used for model training, validation, and testing of ESCAlignNet.

| Total number of individuals | Age in days | Region | Training | Validation | Testing |
| --- | --- | --- | --- | --- | --- |
| 450 | 31--122dpf | lateral body | 12,114 | 3,028 | 500 |
|  |  | dorsal head | 6,793 | 754 | 500 |

Supplementary Table 2 provides detailed distribution statistics of the data used to train, validate, and test the zebrafish feature extraction and matching model (IDNet). The test set comprises three subsets: Test Set 1, which is used to determine the model’s decision threshold and evaluate the model under the optimal threshold; Test Set 2, which is used to test long-term validity; and Test Set 3, which is used to test a short-term photographed cohort of 300 individuals.

Supplementary Table 2. Dataset sizes used for model training, validation, and testing of IDNet. The lateral body and dorsal head are designated as "body" and "back" in the Table.

| Model Stages | Number of individuals used | Region | Training | Validation | Test Set 1 | Test Set 2 | Test Set 3 |
| --- | --- | --- | --- | --- | --- | --- | --- |
| Stage 1 | 408 | Body-Left | 26,371 | 266 | 6,000 | 33,637 | 3,000 |
|  | 408 | Body-Right | 24,833 | 248 | 6,000 | 31,982 | 3,000 |
|  | 434 | Back | 6,962 | 72 | 6,000 | 37,735 | 3,000 |
| Stage 2 | 441 | Body-Left | 34,923 | 350 | 6,000 | 48,660 | 3,000 |
|  | 441 | Body-Right | 30,939 | 310 | 6,000 | 44,489 | 3,000 |
|  | 432 | Back | 5,765 | 58 | 6,000 | 43,394 | 3,000 |

Supplementary Table 3 presents test results for ESCAlignNet. ESCAlignNet uses a dual-branch architecture. To enable comparative testing with the original YOLOv8, the evaluations are divided into two categories: segmentation and keypoint recognition.

Supplementary Table 3. Test results of ESCAlignNet (the improved YOLOv8).

| Models | mAP@0.5  (Mask) | mAP@0.5  (Pose) | mAP@0.5:0.95  (Mask) | mAP@0.5:0.95  (Pose) | Precision | Recall | FPS | Layer |
| --- | --- | --- | --- | --- | --- | --- | --- | --- |
| YOLOv8-seg | 0.968 | -- | 0.782 | -- | 0.940 | 0.922 | 240 | 261 |
| YOLOv8-Pose | -- | 0.955 | -- | 0.720 | 0.924 | 0.930 | **241** | **261** |
| ESCAlignNet-seg | **0.988** | -- | 0.867 | -- | **0.970** | **0.968** | 106 | 377 |
| ESCAlignNet-Pose | -- | **0.970** | -- | 0.811 | **0.977** | **0.955** | 110 | 377 |

Supplementary Table 4 shows the changes in average SSIM values for 100 images per region (dorsal head and lateral body) for three randomly selected individuals before and after segmentation and alignment standardization.

Supplementary Table 4. SSIM statistics of images from different regions and individuals before and after segmentation and alignment standardization. The lateral body and dorsal head are designated as "body" and "back" in the Table.

|  | Zebrafish ID | Body-Left | Body-Right | Back |
| --- | --- | --- | --- | --- |
| SSIM evaluation of 100-image dataset (Before) | 1 | 0.5017 | 0.5062 | 0.3885 |
|  | 2 | 0.4953 | 0.4918 | 0.3917 |
|  | 3 | 0.5446 | 0.5066 | 0.4422 |
|  | Mean | 0.5138 | 0.5016 | 0.4075 |
|  | Median | 0.5117 | 0.4963 | 0.4086 |
| SSIM evaluation of 100-image dataset (After) | 1 | 0.5498 | 0.5534 | 0.5903 |
|  | 2 | 0.5957 | 0.5229 | 0.4825 |
|  | 3 | 0.5722 | 0.5806 | 0.5998 |
|  | Mean | 0.5726 | 0.5523 | 0.5576 |
|  | Median | 0.5463 | 0.5537 | 0.5823 |

Supplementary Table 5 shows the results of Test Set 1 for the IDNet model, which was trained using Stage 1 lateral body data and different combinations of backbone networks and loss functions. All test data underwent data standardization.

Supplementary Table 5. Test results on Test Set 1 for Stage 1 IDNet (trained based on lateral body data) model with different backbone network and loss function combinations.

| Backbone | Loss function | AUC | Validation rate | Accuracy | Best thresholds | FPS | Weights/MB | Data standardization（Yes or No） |
| --- | --- | --- | --- | --- | --- | --- | --- | --- |
| Mobilenet | Triplet loss | 0.860 | 0.7001 | 0.7688 | 1.10 | 218 | 16.2 | Yes |
| Inception_Resnetv1 | Triplet loss | 0.90 | 0.785 | 0.8029 | 1.14 | 168 | 87.3 | Yes |
| IResnet50 | AAM loss | 0.95 | 0.8903 | 0.955 | 1.10 | 122 | 166 | Yes |
| IResnet100 | AAM loss | 0.98 | 0.9406 | 0.960 | 0.99 | 68 | 249 | Yes |
|  | Dynamic AAM loss | **0.99** | **0.9500** | 0.965 | 1.00 | 66 | 249 | Yes |
| IResnet200 | AAM loss | **0.99** | 0.8847 | 0.980 | 0.99 | 35 | 454 | Yes |
|  | Dynamic AAM loss | **0.99** | 0.9203 | **0.98167** | 1.00 | 35 | 454 | Yes |

Supplementary Table 6 shows the results of Test Set 1 for the IDNet model, which was trained using Stage 1 dorsal head data and different combinations of backbone networks and loss functions. Some of the test data did not undergo data standardization.

Supplementary Table 6. Test results on Test Set 1 for Stage 1 IDNet (trained based on dorsal head data) model with different backbone network and loss function combinations.

| Backbone | Loss Function | AUC | Validation rate | Accuracy | Best thresholds | FPS | Weights/MB | Data standardization（Yes or No） |
| --- | --- | --- | --- | --- | --- | --- | --- | --- |
| Inception_Resnetv1 | Triplet loss | 0.705 | 0.680 | 0.608 | 1.10 | 171 | 87.3 | No |
|  | Triplet loss | 0.802 | 0.708 | 0.798 | 1.10 | 172 | 87.3 | Yes |
| Mobilenetv1 | AAM loss | 0.870 | 0.76967 | 0.9072 | 1.19 | 201 | 17.3 | Yes |
| IResnet50 | AAM loss | 0.8777 | 0.68567 | 0.9222 | 1.20 | 67 | 249 | Yes |
| IResnet200 | AAM loss | 0.8755 | 0.56976 | 0.92767 | 1.19 | 35 | 454 | Yes |
|  | Dynamic AAM loss | 0.900 | 0.50667 | 0.9008 | 0.90 | 32 | 454 | Yes |
| IResnet100 | AAM loss | 0.9008 | **0.801** | 0.9333 | 1.22 | 65 | 249 | Yes |
|  | Dynamic AAM loss | **0.950** | 0.7203 | **0.9430** | 1.02 | 63 | 249 | Yes |

Supplementary Table shows the results of Test Set 1 for the IDNet model, which was trained using Stage 2 lateral body data and different combinations of backbone networks and loss functions. All test data underwent data standardization.

Supplementary 7. Test results on Test Set 1 for Stage 2 IDNet (trained based on lateral body data) model with different backbone network and loss function combinations.

| Backbone | Loss function | AUC | Validation rate | Accuracy | Best thresholds | FPS | Weights/MB | Data standardization（Yes or No） |
| --- | --- | --- | --- | --- | --- | --- | --- | --- |
| Mobilenetv1 | Triplet loss | 0.90 | 0.702 | 0.876 | 1.10 | 218 | 16.2 | Yes |
|  | AAM loss | 0.95 | 0.886 | 0.903 | 1.10 | 201 | 17.3 | Yes |
| Inception_Resnetv1 | Triplet loss | 0.95 | 0.786 | 0.886 | 1.14 | 168 | 87.3 | Yes |
| IResnet50 | AAM loss | 0.95 | 0.917 | 0.965 | 1.10 | 122 | 166 | Yes |
| IResnet100 | AAM loss | 0.98 | 0.940 | 0.973 | 0.99 | 68 | 249 | Yes |
|  | Dynamic AAM loss | **0.99** | 0.940 | 0.979 | 1.00 | 66 | 249 | Yes |
| IResnet200 | AAM loss | **0.99** | 0.952 | 0.981 | 0.99 | 35 | 454 | Yes |
|  | Dynamic AAM loss | **0.99** | **0.962** | **0.988** | 1.00 | 35 | 454 | Yes |

Supplementary Table 8 shows the results of Test Set 1 for the IDNet model, which was trained using Stage 2 dorsal head data and different combinations of backbone networks and loss functions. Some of the test data did not undergo data standardization.

Supplementary 8. Test results on Test Set 1 for Stage 2 IDNet (trained based on dorsal head data) model with different backbone network and loss function combinations.

| Backbone | Loss Function | AUC | Validation rate | Accuracy | Best thresholds | FPS | Weights/MB | Data standardization（Yes or No） |
| --- | --- | --- | --- | --- | --- | --- | --- | --- |
| IResnet200 | Dynamic AAM loss | 0.69 | 0.660 | 0.980 | 1.01 | 32 | 454 | Yes |
| IResnet200 | AAM loss | 0.79 | 0.860 | 0.978 | 1.02 | 35 | 454 | Yes |
| Inception_Resnetv1 | Triplet loss | 0.79 | 0.586 | 0.762 | 1.10 | 171 | 87.3 | No |
| Inception_Resnetv1 | Triplet loss | 0.87 | 0.620 | 0.801 | 1.10 | 172 | 87.3 | Yes |
| IResnet100 | AAM loss | 0.900 | 0.902 | 0.913 | 1.02 | 67 | 249 | No |
| IResnet100 | AAM loss | 0.98 | 0.933 | **0.972** | 1.02 | 65 | 249 | Yes |
| IResnet100 | Dynamic AAM loss | **0.99** | **0.939** | **0.972** | 1.01 | 63 | 249 | Yes |
